# The lipid raft marker flotillin FloA drives relocalization of the plasma membrane H^+^-ATPase PmaA as a protective response to calcium stress

**DOI:** 10.64898/2026.04.27.721028

**Authors:** Moemi Kawashima, Thomas Krüger, Maira Rosin, Sophie Tröger-Görler, Thorsten Heinekamp, Axel A. Brakhage

## Abstract

Biological membranes are laterally heterogeneous and contain specialized microdomains called lipid rafts. Lipid rafts serve as organizational platforms that cluster signaling molecules or modulate membrane protein conformation through their unique lipid environment. There are specific lipid raft marker proteins whose functions remain obscure. One of these proteins is flotillin which has been linked to endocytosis. Here, we investigated the regulation and function of FloA, the sole flotillin homolog in the model fungus *Aspergillus nidulans*. FloA expression is specifically upregulated in response to calcium stress, which is a regulatory pattern also conserved in *Aspergillus fumigatus*. Whereas in *A. fumigatus floA* is regulated by the calcium regulatory protein CrzA, this is not the case in *A. nidulans*. BioID proximity labeling revealed that *A. nidulans* FloA physically interacts with proteins in the endocytic pathway as well as another lipid raft marker, the plasma membrane H^+^-ATPase PmaA. Under calcium stress, PmaA undergoes internalization from the cytoplasmic membrane. However, when *floA* is deleted, PmaA internalization is prevented, resulting in cell death. Together, we demonstrate that FloA is essential for the internalization of PmaA during calcium stress, a process that prevents intracellular calcium overload and promotes cell viability. Our results also provide further evidence for flotillin-assisted endocytosis.

**Author abstract:** Lipids and proteins in a cell membrane can cluster together in small regions often called “lipid rafts”, which help the cell interact with its surroundings. Lipid rafts can bring receptors together or influence how membrane proteins behave. Flotillin is a protein which is often found within lipid rafts, but its exact role is not well understood. Instead of using complex mammalian systems, we studied flotillins in the fungus *Aspergillus nidulans*, which is a simpler model organism that allows for a better understanding of cellular processes.

We found that more flotillins are produced when the fungus is exposed to calcium stress. When flotillins were missing, the cells were unable to remove the protein PmaA from the cell membrane during calcium stress. As a result, the fungus could not cope with the calcium stress and eventually died. Therefore, we propose that flotillins are important for the fungus to reorganize its membranes and coping with calcium stress.

## Introduction

Despite biological membranes originally being thought to be uniform mixtures of lipids and proteins, numerous studies have shown that membranes are laterally heterogeneous and contain specialized microdomains called lipid rafts [1–7]. These domains are enriched in sterols and sphingolipids and exhibit both spatial and temporal dynamics. Lipid rafts serve as organizational platforms that cluster signaling molecules or modulate membrane protein conformation through their unique lipid environment [1, 7, 8]. They are implicated in essential cellular processes, including membrane trafficking [9–11], immune signaling [12, 13] and host–pathogen interactions [14–16], and are associated with various human diseases, such as cancer [17–20], cardiovascular disorders [21–23] and susceptibility to infection [14]. Despite their biological significance, lipid rafts remain poorly understood due to their nanometer-scale dimensions, which hinder their direct detection in living cells.

While the complete proteomic composition of lipid rafts remains undefined, members of the stomatin/prohibitin/flotillin/HflK/C (SPFH) domain-containing protein family are consistently identified as raft-associated components across organisms ranging from bacteria to mammals. Among them, flotillins, also known as reggies, are the most extensively characterized. Flotillins show dynamic subcellular localization that varies with cell type and are generally found at the cytoplasmic membrane, late and recycling endosomes, phagosomes, and exosomes [24]. Numerous studies have linked flotillins to endocytosis, suggesting they function in clathrin- and caveolin-independent endocytosis [24–28]. More recent findings, however, propose that flotillins may also function upstream of endocytosis by organizing and clustering cargo molecules at the cytoplasmic membrane before internalization [29].

To gain further insight into flotillin function, we used the filamentous fungus *Aspergillus nidulans* as a eukaryotic model system. Compared to mammalian cells, this organism is easier to handle and provides a genetically modifiable model for studying membrane dynamics in a multicellular eukaryote. In contrast to mammals, which express two flotillin paralogs, *A. nidulans* encodes a single homolog, *floA* [30], simplifying functional analysis. We demonstrate that *floA* expression is specifically induced under calcium stress and that FloA is essential for the internalization of the plasma membrane H^+^-ATPase PmaA under these conditions. This internalization prevents intracellular calcium overload and is critical for cell survival. Together, our results establish a mechanistic link between flotillins and the calcium stress response, placing flotillins in a new functional context and providing further evidence for flotillin-assisted endocytosis.

## Results

### FloA localizes to the plasma membrane

In order to characterize the biological function of FloA, a *floA* knockout mutant was generated. The knockout was confirmed by PCR (S4 Fig.). The *floA* knockout mutant showed no difference in colony diameter compared to the wild-type strain R21 on AMM agar plates and showed no obvious morphological defects when observed microscopically (Fig. 1a). These results indicate that *floA* is a gene that is not required for the normal growth of *A. nidulans* under standard laboratory conditions. Furthermore, FloA was tagged with GFP at its C-terminus and expressed under the control of the inducible *alcA* promoter (*alcA*_p_), as the native promoter is too weak for microscopic observation [30]. FloA-GFP signals were observed as punctate structures on the plasma membrane, with the strongest signals appearing in conidia (Fig. 1b).

**Figure 1.**
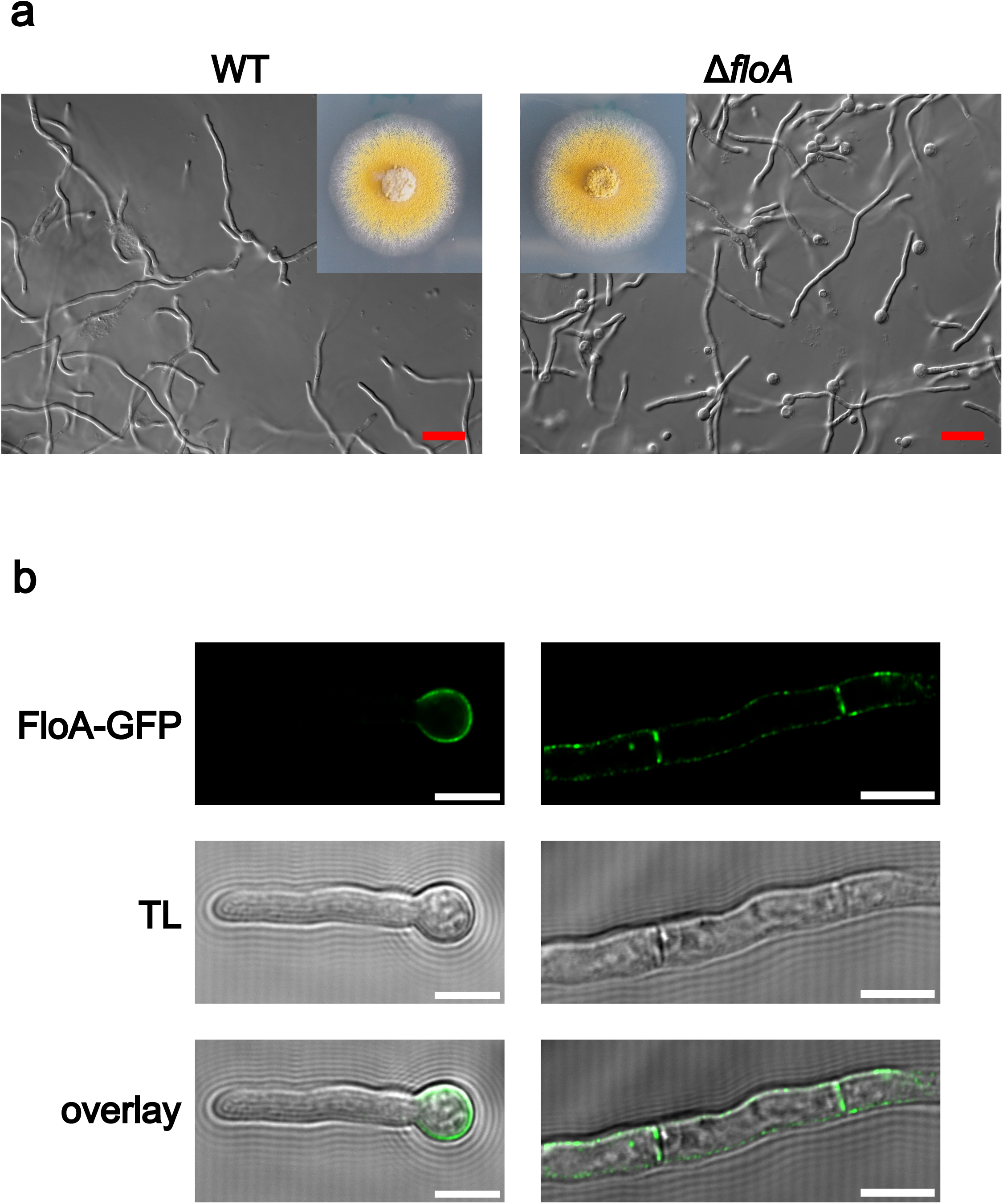
Characterization of *floA* deletion strains and FloA localization. (a) Growth comparison of wild-type and Δ*floA* strain. Scale bar 20 µm. (b) Localization of FloA-GFP produced under the control of *alcA*_p_. To image FloA-GFP in hyphae, acquisition settings were adjusted to capture weaker signals. Scale bar 5 µm.

### FloA expression is specifically induced by calcium stress

The absence of an obvious phenotype in the *floA* knockout mutant under standard growth conditions suggested that FloA is required only under specific physiological states. To determine the conditions under which FloA becomes functionally relevant, its expression was quantified under various stress conditions using a nanoluciferase (nLuc) fusion construct (Fig. 2a). The abundance of FloA was extremely low under the non-stress conditions, again supporting the notion that the protein is not required during normal growth. In contrast, FloA expression strongly increased in a dose-dependent manner upon exposure to calcium stress. While upregulation was moderate in 1 mM CaCl_2_, the combination of 1 mM CaCl_2_ with the calcium ionophore A23187 resulted in upregulation comparable to that observed in 0.2 M CaCl_2_, indicating that the elevated intracellular calcium level is the primary trigger. This induction was specific to calcium stress and not observed in response to other ionic or osmotic stressors, namely 0.2 M MgCl_2_, 0.2 M KCl, 0.2 M NaCl and 0.6 M sorbitol. The induction by calcium was also observed microscopically using a strain expressing FloA-GFP under the native *floA* promoter (Fig 1b).

**Figure 2.**
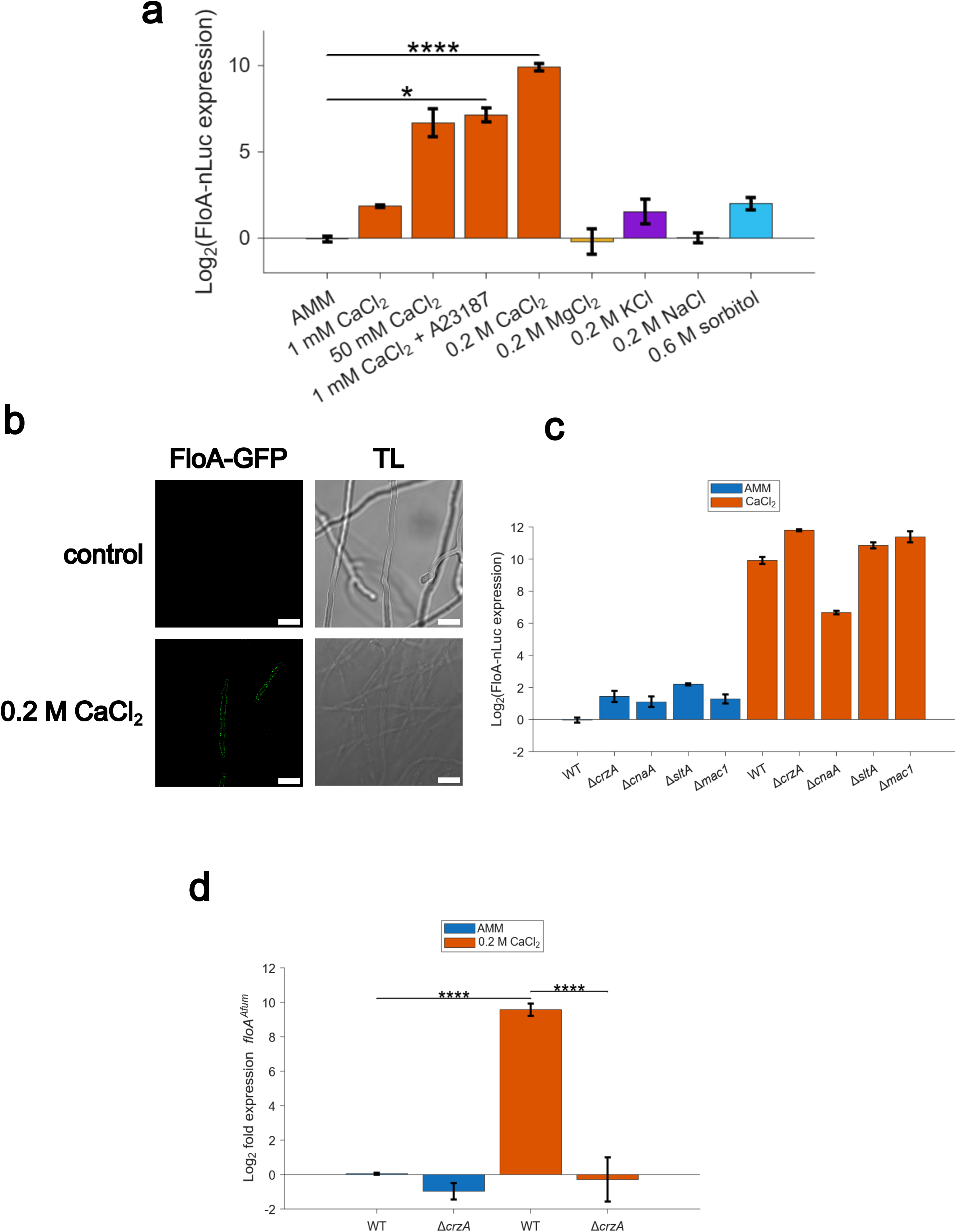
Induction of *A. nidulans* FloA by calcium stress without involvement of known calcium-responsive transcription factors. (a) FloA-nLuc abundance under different conditions quantified by luminescence intensity normalized by total protein concentration. (b) FloA-GFP expression in AMM and 0.2 M CaCl_2_. The expression is controlled by the native *floA* promoter. (c) Normalized luminescence intensity of the FloA-nLuc reporter strain in the wild-type and calcium-responsive regulator knockout backgrounds. All strains were cultured in AMM with 0.2 M CaCl_2_, except the Δ*cnaA* strain, which was cultured in 1 mM CaCl_2_ instead of 0.2 M CaCl_2_ due to its strong sensitivity to calcium stress. (d) qRT-PCR analysis indicated as relative *floA* expression in *A. fumigatus* wild type and Δ*crzA* strains. Error bars represent the mean *±* SD. * p *≤* 0.05, ****p *≤* 0.0001. Statistical significance between samples were evaluated by one-way ANOVA followed by Tukey’s multiple comparisons test (a and c).

As CrzA is a major regulator of calcium responses in fungi, we investigated whether *floA* is regulated by this transcription factor. To this end, *crzA* was knocked out in the FloA-nLuc reporter strain and the FloA expression was assessed under 0.2 M CaCl_2_ (Fig. 2c). However, the deletion of *crzA* did not influence *floA* expression, indicating that *floA* is not regulated by CrzA, despite being calcium induced. Furthermore, other known calcium-responsive regulatory genes were knocked out, namely the calcium-sensing protein phosphatase calcineurin, the cation homeostasis-responsive factor *sltA* [31] and the copper starvation-responsive transcription factor *mac1* [32]. However, none of these knockouts abolished the calcium-induced upregulation of FloA.

Interestingly, qRT-PCR analysis confirmed that the homolog of *floA* in *A. fumigatus* was also strongly upregulated under calcium stress in the wild-type strain but not in the *crzA* knockout strain (Fig. 2d). This observation suggests that although calcium-dependent regulation of flotillins may be conserved across *Aspergillus* species, there appears to be a species-specific regulatory divergence between *A. nidulans* and *A. fumigatus*.

### Proximity-dependent biotin identification (BioID) assay indicates the physical interaction of FloA with proteins of the endocytic pathway

Flotillins are proposed to function as scaffolds that facilitate the clustering of other proteins within lipid rafts, rather than exerting intrinsic enzymatic activity [33]. However, the identity of their interaction partners remains largely unknown. To address this, we performed a BioID assay to identify proteins in proximity to FloA. For this purpose, we generated strains expressing either a FloA-3*×*hemagglutinin tag (HA)-miniTurboID fusion protein or unfused 3*×*HA-miniTurboID, the latter serving as a negative control. Expression of both constructs was placed under control of the inducible *alcA* promoter to increase the otherwise low basal expression of FloA and simultaneously minimize background biotinylation, a known limitation of BioID approaches [34, 35]. The expression of 3*×*HA-miniTurboID and FloA-3*×*HA-miniTurboID, and its biotinylation activities were confirmed by Western blot analyses (Fig. 3a).

**Figure 3.**
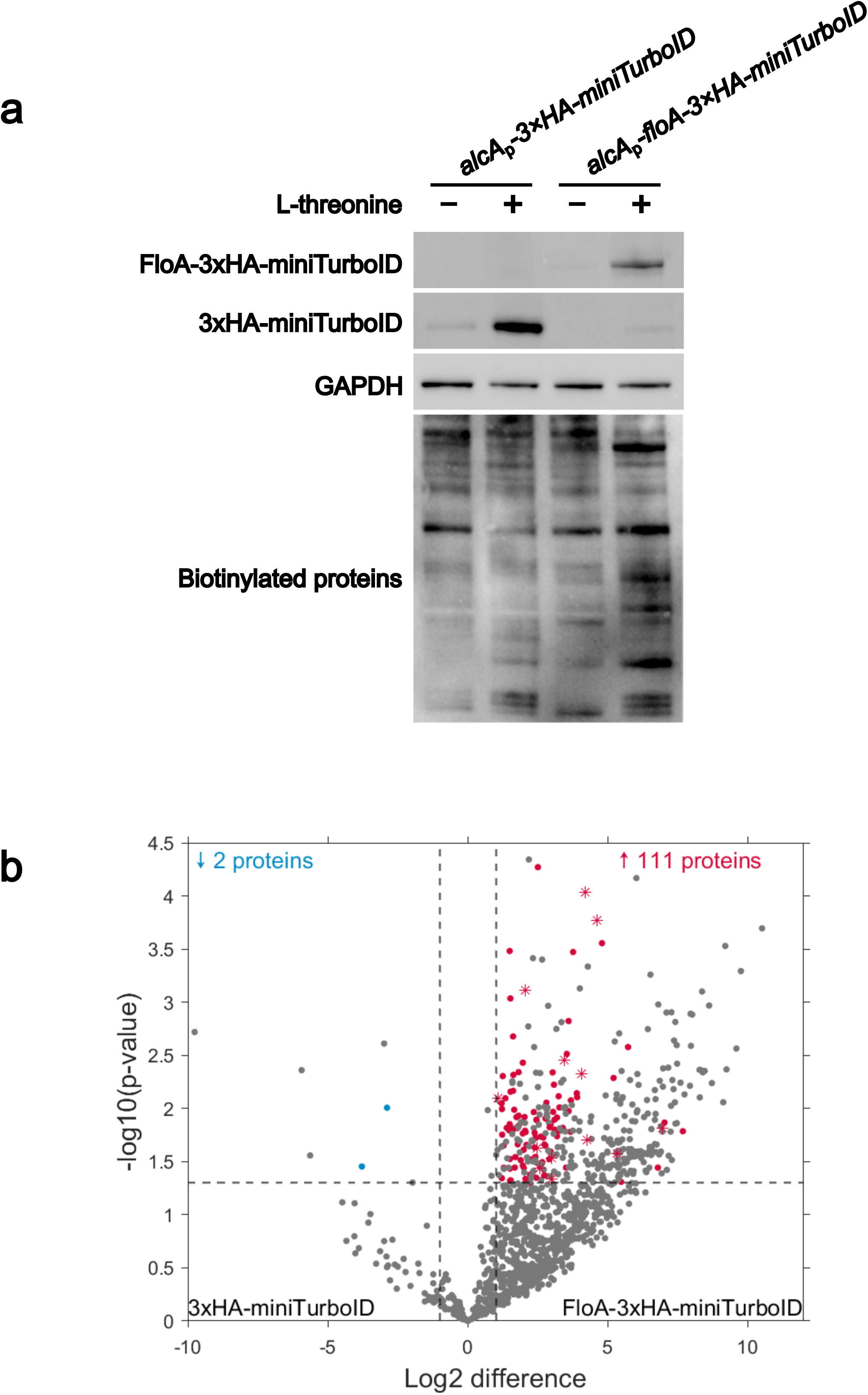
BioID analysis for identification of proteins potentially interacting with FloA. (a) Western blot analyses of 3*×*HA-miniTurboID and FloA-3*×*HA-miniTurboID expressed under the control of *alcA*_p_ under inducing (+ L-threonine) and repressing conditions (-L-threonine). The biotinylation activities of these proteins were verified by detecting biotinylated proteins using HRP-conjugated streptavidin. GAPDH was used as a control. (b) Volcano plot analysis of the BioID data using FloA-3*×*HA-miniTurboID as bait and 3*×*HA-miniTurboID as a negative control. Proteins with statistically significant enrichment for each strain are indicated in red (FloA-3*×*HA-miniTurboID) and blue (3*×*HA-miniTurboID). Asterisks indicate selected proteins that are relevant for the endocytic pathway.

By applying liquid chromatography-tandem mass spectrometry (LC-MS/MS), a total of 1228 proteins were identified, of which 270 were uniquely detected in the *floA-3xHA-miniTurboID* strain. Among these, 111 proteins were significantly enriched compared to the negative control (Fig. 3b). Notably, many of these enriched proteins are associated with the endocytic pathway, consistent with previous reports suggesting a role of flotillins in endocytosis (Table 1). For example, clathrin and actin are both central components of the canonical endocytic pathway, with clathrin coating the endocytic site and actin supplying mechanical force [36]. In addition, the dynamin-like protein VpsA (ANIA 08023) was also enriched, its yeast homolog Vps1 is believed to pinch off the endosome from the plasma membrane [37]. Myosin V and tubulin are components required for membrane trafficking, with myosin V acting as a motor for vesicles moving along cytoskeletal tracks such as microtubules [38]. The Rab GDP dissociation inhibitor and the GTP-binding protein RhoA are both regulators of membrane trafficking, although they are not direct components of the endocytic machinery. A strong indication for the successful enrichment of flotillin-associated proteins by our approach is the enrichment of known lipid raft-associated proteins, namely subunit A of the V-type proton ATPase [14, 39, 40] and the plasma membrane H^+^-ATPase [41–45]. Our data support the hypothesis that FloA plays a role in endocytosis in *A. nidulans*, similar to its role in other eukaryotes.

**Table 1.** Selected BioID enriched proteins identified by LC-MS/MS.

| No. | Gene ID | log2<br>difference | -log10(*p*-value) | Description |
| --- | --- | --- | --- | --- |
| 1 | ANIA_00463 | 5.327 | 1.573 | Clathrin heavy chain |
| 2 | ANIA_05895 | 4.621 | 3.773 | Rab GDP dissociation inhibitor |
| 3 | ANIA_02050 | 4.272 | 1.702 | Clathrin light chain |
| 4 | ANIA_06177 | 4.199 | 4.036 | FloA |
| 5 | ANIA_08023 | 4.064 | 2.325 | Vacuolar protein sorting-<br>associated protein 1 |
| 6 | ANIA_08862 | 3.465 | 2.452 | Myosin V homolog |
| 7 | ANIA_05740 | 2.989 | 1.339 | GTP-binding protein RhoA |
| 8 | ANIA_06542 | 2.980 | 1.534 | Gamma actin |
| 9 | ANIA_00316 | 2.580 | 1.439 | Alpha tubulin |
| 10 | ANIA_08021 | 2.488 | 1.630 | V-type proton ATPase catalytic<br>subunit A |
| 11 | ANIA_04859 | 2.457 | 3.115 | Plasma membrane H<sup>+</sup>-ATPase<br>PmaA |
| 12 | ANIA_01182 | 1.647 | 2.092 | Beta tubulin |

### FloA and PmaA are present in detergent resistant membranes (DRMs) but are localized in distinct membrane domains

Among the proteins enriched in the BioID dataset, the plasma membrane bound H^+^-ATPase PmaA was further investigated as a potential interaction partner of FloA, as its yeast homolog Pma1 is a well-characterized lipid raft-associated protein [41–45]. Despite some inherent limitations, DRM isolation remains the primary biochemical approach for probing lipid raft association of proteins [46]. In this method, membranes are treated with a non-ionic detergent, typically Triton X-100, at low temperature, which preserves sphingolipid- and sterol-enriched domains while solubilizing the bulk of the membrane.

To assess whether FloA and PmaA have raft affinity in *A. nidulans*, we performed DRM fractionation and Western blot analyses (Fig. 4a). As expected, both FloA and PmaA were detected in low-density and high-density fractions, whereas β-tubulin (TubA) was predominantly present in high-density fractions, indicating that both FloA and PmaA have the capacity to associate with lipid rafts.

**Figure 4.**
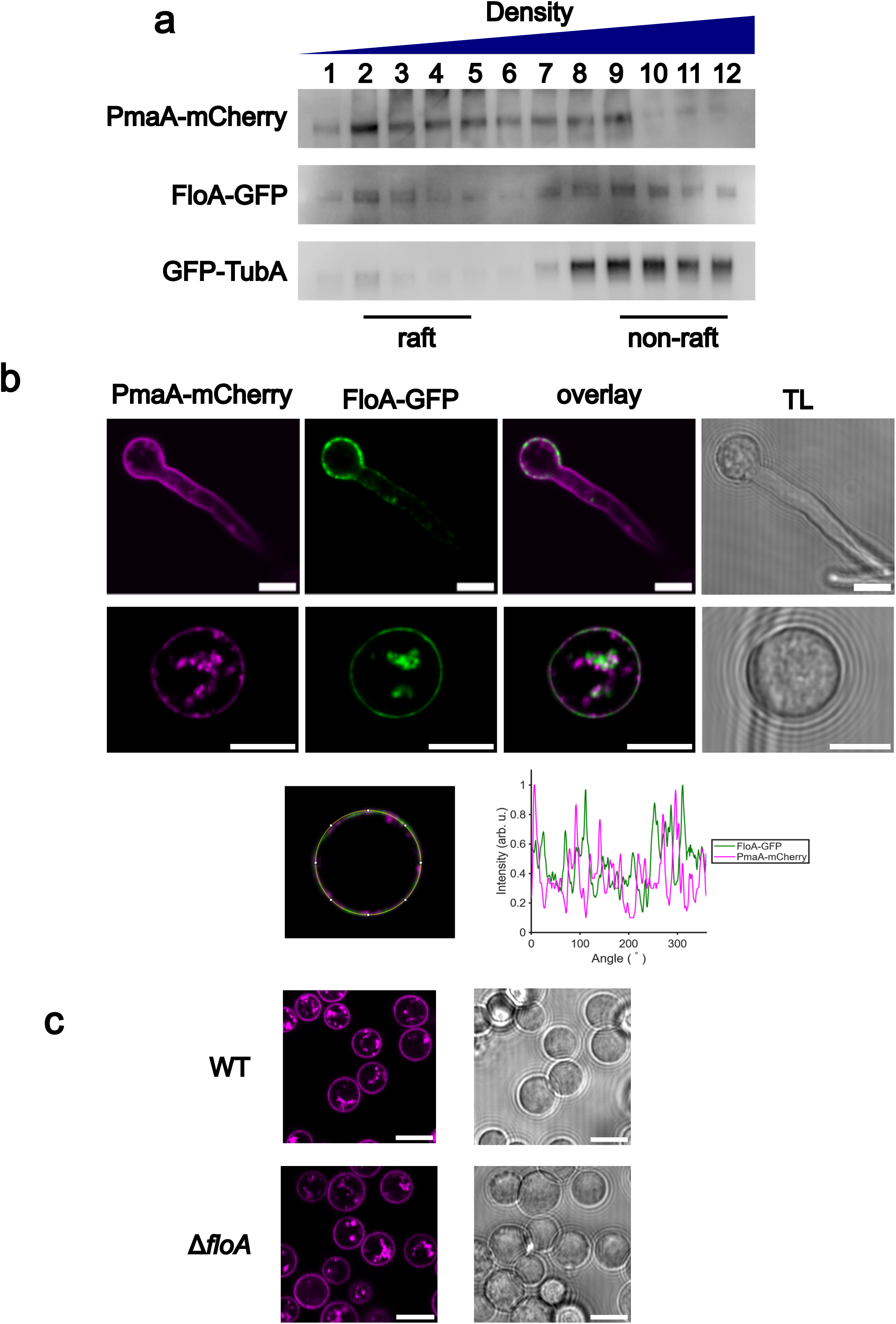
Lipid-raft association and localization of FloA-GFP and PmaA-mCherry. (a) Western blot analysis of FloA-GFP and PmaA-mCherry isolated from DRMs. GFP-TubA was used as a non-raft control. Fusion proteins were detected using anti-GFP and anti-mCherry antibodies. (b) Cellular localization of FloA-GFP and PmaA-mCherry in *A. nidulans* strain *alcA*_p_*-floA-GFP; pmaA-mCherry*. The intensity profile of the two proteins along the plasma membrane of a conidium was plotted to indicate distinct localization. (c) PmaA-mCherry signals in the wild-type and Δ*floA* background. The intensity profiles of the two proteins along the plasma membrane of a conidium were plotted to indicate distinct localization. All scale bars 5 µm.

Lipid rafts are thought to consist of multiple subsets characterized by distinct protein and lipid compositions. To investigate whether FloA and PmaA localize to the same subset, we examined their subcellular distribution by fluorescence microscopy (Fig. 4b). PmaA-mCherry displayed signals primarily on the cytoplasmic membrane as previously reported [47]. Although the signals appeared to be evenly distributed at first glance, airyscan superresolution microscopy revealed that they were clustered in patches. This observation further confirms that PmaA is localized in distinct membrane compartments. The intensity profile of PmaA-mCherry and FloA-GFP along the plasma membrane of a conidium indicated that the punctate structures of the two proteins had little overlap. Furthermore, the deletion of *floA* did not alter the distribution of PmaA-mCherry in the control condition (Fig. 4c). This is in line with the fact that the abundance of FloA is extremely low without high calcium stress and is most likely not an important protein for cellular growth under standard laboratory conditions.

### PmaA is internalized in response to calcium stress

As shown in Fig. 2a, FloA expression is specifically induced by calcium stress. To explore the role of FloA under these conditions, and to assess a potential functional relationship with PmaA, we examined the subcellular localization patterns of FloA-GFP and PmaA-mCherry during calcium stress using fluorescence microscopy (Fig. 5a). The C-terminal tagging of the essential protein PmaA has previously been shown to be non-disruptive [47].

**Figure 5.**
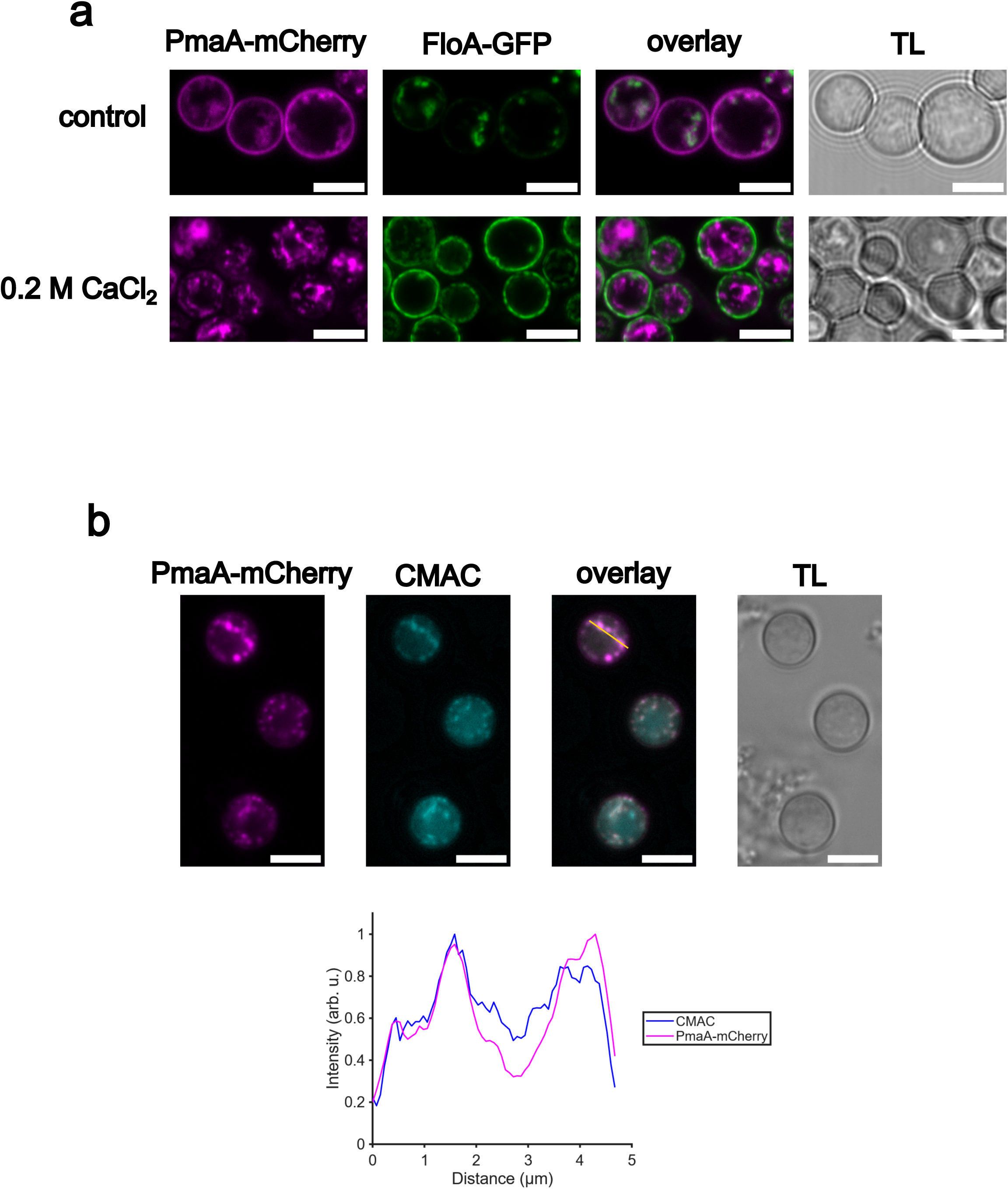
Redistribution of FloA-GFP and PmaA-mCherry under calcium stress. (a) Localization of FloA-GFP and PmaA-mCherry in AMM (control) and 0.2 M CaCl_2_. The conidium was imaged using airyscan superresolution microscopy. (b) Colocalization of PmaA with the late endosomes/vacuole stain CMAC in 0.2 M CaCl_2_. The intensity profiles of PmaA-mCherry and CMAC along the yellow line were plotted to indicate colocalization. All scale bars 5 µm.

In the control condition, FloA-GFP localized to small punctate structures on the cytoplasmic membrane of conidia and showed additional intracellular signals (Fig. 5a). However, upon exposure to calcium stress, FloA-GFP redistributed to form a uniform signal almost exclusively along the cytoplasmic membrane of conidia. In contrast, PmaA-mCherry exhibited the opposite behavior and redistributed from the cytoplasmic membrane to the cytoplasm under calcium stress.

As Pma1 internalization is associated with vacuolar degradation [41], we stained acidic compartments using CMAC (7-amino-4-chloromethylcoumarin) (Fig. 5b). As expected, internalized PmaA-mCherry colocalized with CMAC signals, indicating its localization at late endosomes/vacuoles.

### PmaA internalization requires FloA and promotes cell survival under calcium stress

To assess whether FloA contributes to PmaA internalization during calcium stress, PmaA-mCherry localization in the Δ*floA* background was inspected. In the Δ*floA* background, PmaA-mCherry failed to internalize upon calcium stress (Fig. 6a). As expected, PmaA internalization and the impairment thereof could only be observed in 0.2 M CaCl_2_ but not in 0.2 M MgCl_2_, 0.2 M NaCl, 0.2 M KCl and 0.6 M sorbitol (Fig. 6a and S5 Fig.). This suggests that FloA is only important for PmaA internalization under calcium stress but not for other ionic and osmotic stressors. We then addressed the question whether PmaA internalization contributes to calcium homeostasis and cell viability under calcium stress. Therefore, we employed the calcium indicator Fluo-4 AM together with the membrane-impermeable viability dye Sytox Blue (Fig. 6b). In the wild-type strain, the majority of conidia internalized PmaA-mCherry upon calcium treatment and remained negative for both Fluo-4 and Sytox Blue. Conversely, in the Δ*floA* background, a substantial fraction of conidia failed to internalize PmaA-mCherry. Such cells were positive for Fluo-4 and/or Sytox Blue. The complementation of *floA* restored this phenotype and the complemented strain behaved similarly to the wild-type strain. These findings indicate that PmaA internalization under calcium stress prevents intracellular calcium overload and promotes cell survival. Taken together, the data support a model in which FloA enables PmaA relocalization as part of a protective response to calcium stress.

**Figure 6.**
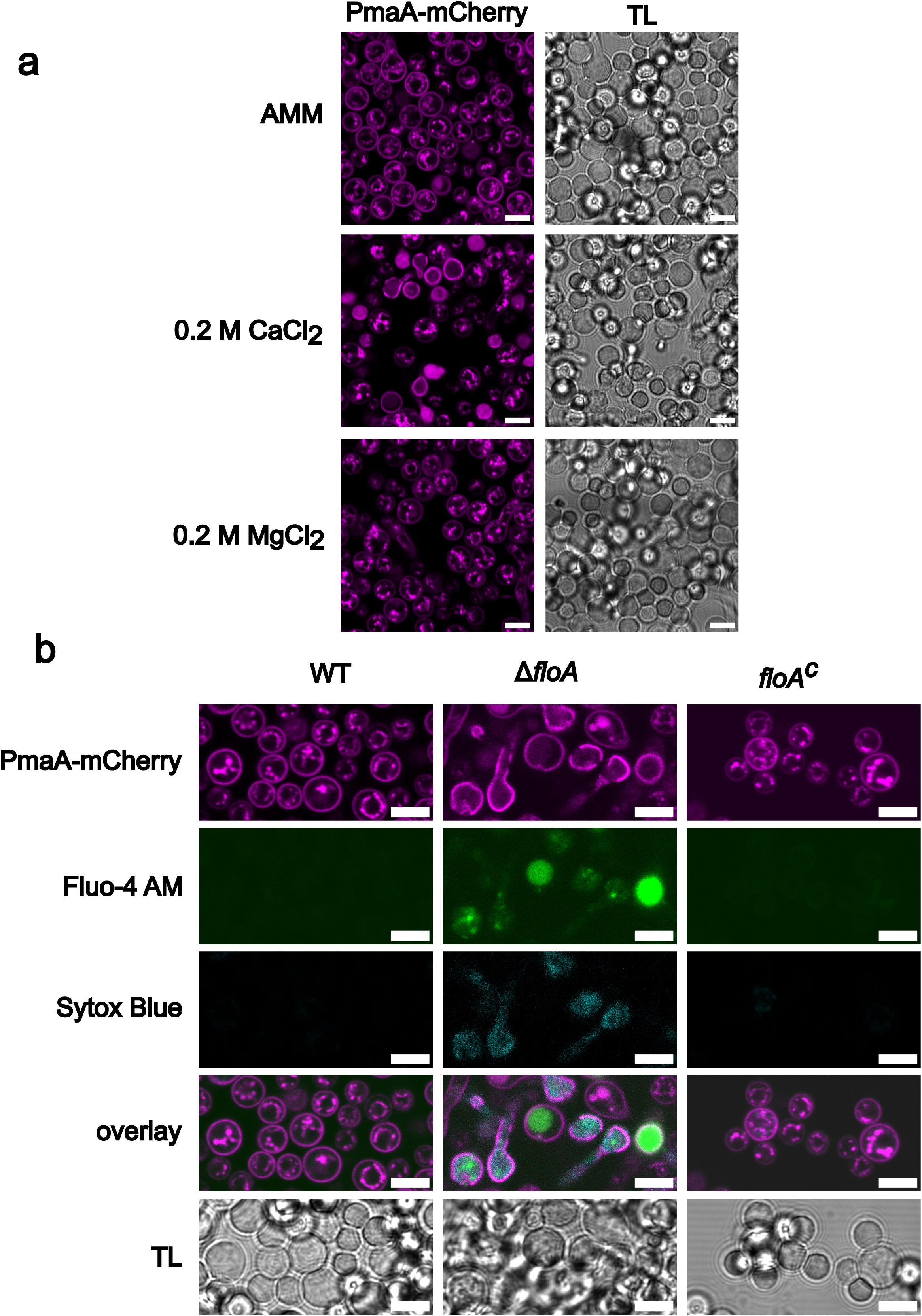
Impact of FloA-dependent PmaA internalization on cell viability. (a) Localization of PmaA-mCherry in the Δ*floA* background in AMM, 0.2 M CaCl_2_ and 0.2 M MgCl_2_. (b) Intracellular calcium levels and cell viability evaluated by Fluo-4 and Sytox Blue, respectively, in the wild type, Δ*floA* and *floA* complementation strains in 0.2 M CaCl_2_. All scale bars 5 µm.

## Discussion

Although early studies on lipid raft focused on mammalian cells, the yeast *Saccharomyces cerevisiae* has also been widely used as a model to investigate raft properties. Yeast and mammalian cells share similar plasma membrane compositions, and many general properties of lipid rafts are conserved [48]. For example, the fungal membrane compartment of Can1 overlaps with eisosomes [49], which share several characteristics with mammalian caveolae [50], a distinct subset of lipid rafts. One major difference, however, is that fungi use ergosterol as their main sterol, which has a stronger raft-forming capacity than cholesterol, the major sterol in mammals [51]. Due to such membrane properties, fungal systems can offer valuable insights into mammalian raft biology.

However, *S. cerevisiae* lacks most proteins of the SPFH family, except for prohibitins [30, 52], limiting its usefulness for studying flotillin biology. In contrast, the model filamentous fungus *A. nidulans* encodes homologs of flotillin, stomatin and prohibitin [30]. Therefore, we performed a detailed functional analysis of FloA in *A. nidulans*, given that flotillins are among the most extensively studied lipid raft marker proteins in mammalian cells.

By tagging FloA with nanoluciferase, we found that FloA expression is specifically induced by calcium stress. Although FloA is barely detectable under non-stress conditions, its expression increased nearly a thousand-fold in response to 0.2 M CaCl_2_, but not to other ionic or osmotic stressors. Thus, we discovered a highly specific calcium-dependent induction of a fungal flotillin. The connection between flotillins and calcium has also been found in other organisms. For example, the plant flotillin Flot1 in *Arabidopsis thaliana* is among the top upregulated genes in response to high humidity, which rapidly increases the cytosolic calcium concentrations in leaves [53]. Flotillins are also highly expressed in neuronal tissues of zebrafish [54] and *Xenopus laevis* [55], and have been linked to neuronal regeneration [56] and the formation of glutamatergic synapses [57], where calcium signalling is of critical importance. In addition, it has been shown that calmodulin activation is likely required for forming flotillin-rich membrane domains on phagolysosomal membranes in macrophages [14].

In some contexts, calcium also regulates flotillins post-translationally. For instance, flotillin-2 in human intestinal epithelial cells exhibits a calcium-dependent relocalization from perinuclear regions to the cytoplasmic membrane upon CaCl_2_ treatment [58]. Taken together, these data suggest that flotillins play important roles in processes in which intracellular calcium levels need to be tightly regulated and that calcium serves as a conserved spatial regulator of flotillins. This could explain why FloA-GFP signals accumulate in *A. nidulans* conidia, since conidia must survive very harsh conditions and therefore may require strict regulation of intracellular calcium levels.

Signal transduction in eukaryotes often involves the canonical calcium-calmodulin-calcineurin signaling pathway[59]. When the intracellular calcium level rises, calmodulin binds Ca^2+^ and subsequently activates the serine/threonine phosphatase calcineurin, which then dephosphorylates downstream targets, which in fungi include the main calcium-responsive transcription factor Crz1/CrzA [60]. Although FloA expression was induced by elevated intracellular calcium, surprisingly, our data revealed that this induction was independent of CrzA, as well as calcineurin and other known calcium-responsive transcription factors. Interestingly, however, the expression of the FloA homolog in the distantly related human-pathogenic species *A. fumigatus* was shown to be upregulated under calcium stress in a CrzA-dependent manner. Such species-specific differences in the calcineurin pathway have also been observed in other fungi. For example, calcineurin is essential for growth at 37 °C in *Cryptococcus neoformans* [61–63], but not in *Candida albicans* [64], although it is required for virulence in both species. Our findings support the hypothesis that, although the calcineurin pathway is conserved across fungi, its downstream targets and physiological importance can vary between species [65].

Flotillins are generally viewed as scaffolding proteins that organize other molecules within lipid raft microdomains, rather than carrying out enzymatic functions themselves [33]. To understand which proteins might be recruited to lipid rafts with the help of flotillins, we carried out a BioID assay to map proteins that interact with FloA or are part of common complexes. The dataset revealed several components of the endocytic machinery, including clathrin, myosin and actin. Another interesting hit was tubulin, which forms the microtubule used for endosomal transport. These findings agree with the hypothesis that flotillins are linked to endocytosis. Moreover, our analysis identified well-known raft-associated proteins, including subunits of the vacuolar H^+^-ATPase [14, 39, 40] and the plasma membrane H^+^-ATPase [41–45], which further validates our experimental approach.

PmaA is an essential protein that pumps protons out of the cell to regulate cellular pH and to create an electrochemical gradient across the plasma membrane [66, 67]. In *S. cerevisiae*, Pma1 is synthesized in the endoplasmic reticulum (ER) and transported to the cytoplasmic membrane *via* the Golgi apparatus, where it associates with lipid rafts prior to surface delivery [68–70]. The *A. nidulans* PmaA protein shares 50% similarity with Pma1 [71], yet follows an unconventional trafficking pathway, bypassing the Golgi apparatus after its exit from the ER [47]. In *S. cerevisiae*, Pma1 interacts with Ast1, which promotes its recruitment to lipid rafts and prevents its mis-sorting to the vacuole [42, 72]. However, potential interactions between Pma1 and flotillins have not been examined in the yeast model, as flotillins are absent from *S. cerevisiae*. Here, we investigated whether *A. nidulans* PmaA associates with lipid rafts and tested whether FloA, similar to Ast1, can support PmaA recruitment to raft domains. Density gradient centrifugation demonstrated that both PmaA and FloA are present in DRMs, supporting their association with lipid raft domains. Although PmaA-mCherry and FloA-GFP localized in the plasma membrane under standard conditions, they did not colocalize with each other, implying that they reside in distinct subsets of membrane microdomains. Under these conditions, PmaA-mCherry was mainly found at the cytoplasmic membrane with only weak intracellular signals, while FloA-GFP showed the opposite pattern.

The downregulation of membrane proteins by endocytic internalization is a well-established regulatory mechanism, typically followed by their sorting to lysosomes or vacuoles for degradation [73–76]. Here, we demonstrate that PmaA undergoes internalization in response to calcium stress. Upon exposure to 0.2 M CaCl_2_, the distribution of FloA and PmaA shifted markedly: PmaA-mCherry became strongly internalized into late endosomes and/or vacuoles, while FloA-GFP accumulated at the cytoplasmic membrane. Strikingly, in the Δ*floA* background, PmaA-mCherry failed to undergo calcium-induced internalization, indicating a mechanistic role of FloA for the endocytic process. Loss of FloA resulted in uncontrolled calcium uptake and subsequent cell death.

Calcium enters fungal cells mainly through the voltage-gated Cch1/CchA channel and the stretch-activated Mid1/MidA channel [77]. After entry, calcium is primarily sequestered into the vacuole *via* Ca^2+^-ATPases and the Ca^2+^/H^+^ exchanger [78–80]. V-ATPase activity is essential for calcium homeostasis, as it generates the proton gradient required for Ca^2+^/H^+^ exchange [78]. Our findings uncovered that the lack of PmaA internalization led to cell death under calcium stress. There are two non-mutually exclusive explanations for the downstream consequences of PmaA endocytosis. First, removing PmaA from the cytoplasmic membrane reduces proton pumping at the cell surface, causing membrane depolarization and thereby lowering the activity of voltage-gated Ca^2+^ channels. In *S. cerevisiae*, Ca^2+^ influx is mediated by channels that open in response to membrane hyperpolarization [81]. Moreover, reduced Pma1 activity has been reported to decrease Cch1/Mid1 channel activity [82]. Second, if internalized PmaA remained active on endosomal membranes, it pumps protons into endosomes, increasing the proton gradient that drives the Ca^2+^/H^+^ exchanger and thus speeding up vacuolar calcium sequestration. Previous studies have shown that the loss of V-ATPase activity in *S. cerevisiae* results in ubiquitination and endocytic downregulation of Pma1 [83–85], enabling the protein to reach endosomal compartments where it may partially compensate for the absence of V-ATPase-driven acidification [85]. Furthermore, the endocytic uptake of Pma1 in V-ATPase deficient mutants requires calcineurin activation [85]. Both proposed mechanisms would help prevent cytosolic calcium overload. However, further experiments are required to determine whether PmaA internalization truly alters the cytoplasmic membrane potential and whether the internalized protein retains proton-pumping activity at endosomes.

Our results also reveal a mechanistic connection between FloA and the endocytic uptake of PmaA under calcium stress. In *A. thaliana*, flotillins regulate cytoplasmic membrane H^+^-ATPase abundance under salt stress by modulating the rate of exocytic delivery rather than influencing endocytosis [86]. Similarly, studies in human cells show that caveolin-1 and flotillin-1/2 can negatively regulate the internalization of membrane proteins [87, 88]. Although our data suggest that FloA is required for PmaA internalization, FloA-GFP accumulated at the cytoplasmic membrane under calcium stress and was not internalized together with PmaA-mCherry after uptake. This suggests that PmaA is not sorted into FloA-containing raft domains on endosomes. Instead, consistent with previously proposed models [29], FloA may act at the cytoplasmic membrane by mediating pre-endocytic clustering of PmaA rather than participating directly in the endocytic machinery (Fig. 7).

**Figure 7.**
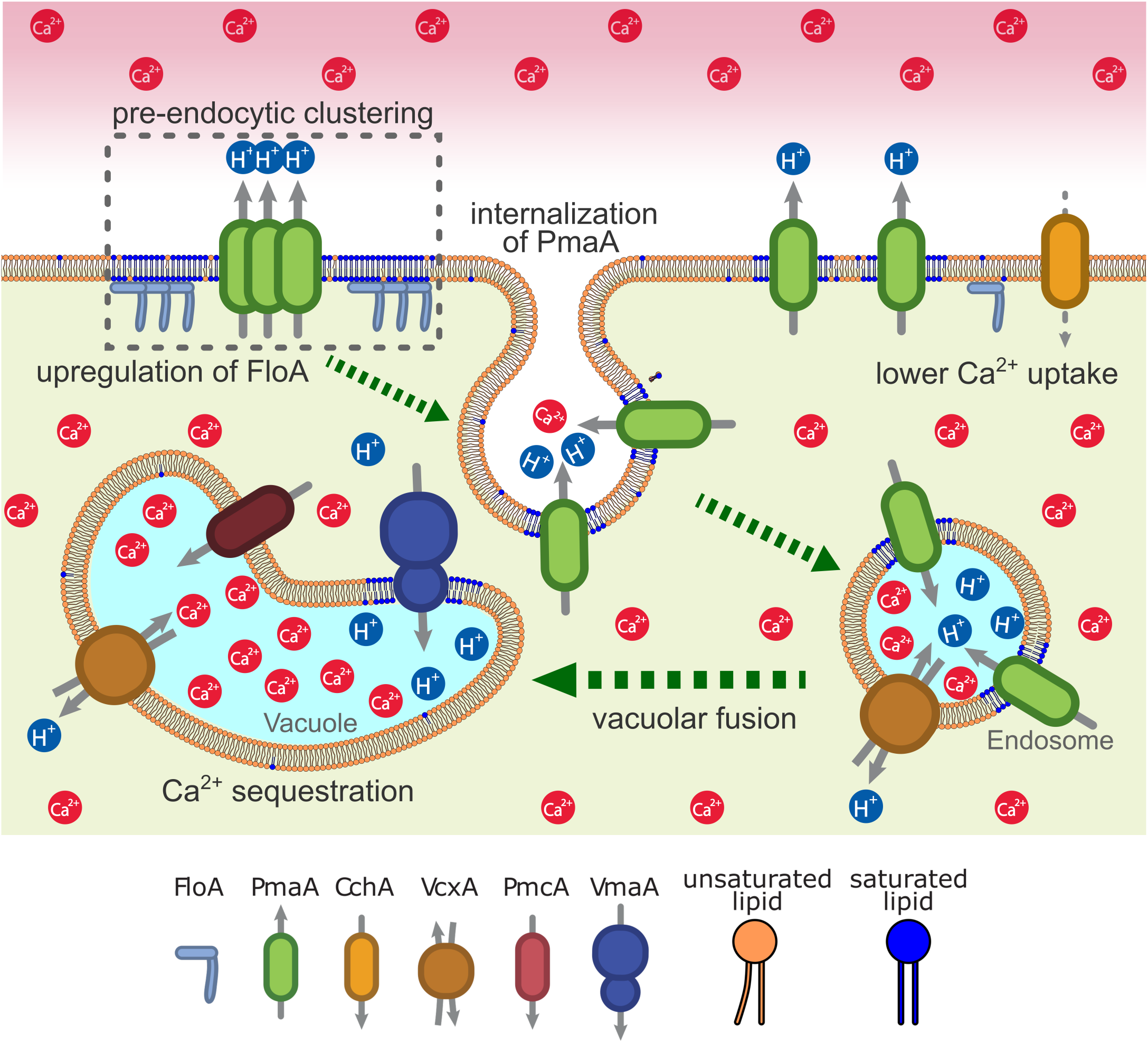
Model of FloA-dependent PmaA internalization under calcium stress. When intracellular calcium levels are elevated, *A. nidulans* responds by strongly upregulating the expression of FloA and reducing the number of PmaA molecules in the cytoplasmic membrane, most likely by endocytosis. FloA contributes to the clustering of PmaA prior to internalization, potentially leading to depolarization of the cytoplasmic membrane and thereby a reduced Ca^2+^ uptake due to lowered Ca^2+^ channel activity. After internalization, PmaA is redistributed at late endosomes/vacuoles. Here, another possible downstream effect is the activation of Ca^2+^/H^+^ exchangers through the proton pumping activity of PmaA on endosomal membranes. This may explain why PmaA internalization is important for Ca^2+^ sequestration and cell survival under calcium stress.

In conclusion, we have shown that FloA is required for PmaA internalization under calcium stress, a process that promotes cell survival. Overall, our study proves *A. nidulans* to be a powerful system for dissecting the functions of SPFH proteins in eukaryotic cells and may ultimately contribute to antifungal strategies targeting the essential plasma membrane H^+^-ATPase.

## Materials and Methods

### Strains, plasmids, media and cultivation

All fungal strains and primers used in this study are listed in S1 Table and S2 Table. Detailed methods of the generation of *A. nidulans* mutant strains can be found in S3 Appendix. *A. nidulans* was cultured in *Aspergillus* minimal medium (AMM) as previously described [89]. When required, biotin (0.3 µg/mL), *para*-aminobenzoic acid (15 µg/mL), L-arginine (0.5 mg/mL), uridine (1.2 mg/mL) or uracil (2.2 mg/mL) were added to the medium. For *Aspergillus fumigatus*, AMM was supplemented with Hutner’s trace elements [90]. Where required, the compound A23187 (Sigma-Aldrich) was added at a final concentration of 5 µg/mL from a stock solution of 2 mg/mL in DMSO. For comparison of growth rates between strains, 10^6^ spores suspended in 10 µL H_2_O were spotted onto AMM agar plates and incubated for 3 days at 37 °C.

Plasmids were propagated in *Escherichia coli* DH10β. *E. coli* was grown in LB medium supplemented with 100 µg/mL carbenicillin.

### Luciferase assay

For quantification of FloA-nLuc, mycelium from *A. nidulans* strain *floA-nLuc* was added to a 2 mL screwcap tube, filled with PBS and zirconia beads (Thermo Fisher Scientific), and then homogenized using a FastPrep homogenizer (MP Biomedicals). After centrifugation, the uminescence of the supernatant was analyzed using the Nano-Glo luciferase assay system (Promega) according to the manufacturer’s instructions. The luminescence intensity was measured with the ClarioSTAR microtiter plate reader (BMG LabTech) and then normalized by the total protein concentration determined by a Bradford assay using the Pierce Bradford Plus Protein Assay Kit (Thermo Fisher Scientific), where the absorbance at 595 nm was measured in three technical replicates using the microtiter plate reader. The standard curve was fitted with a second-order polynomial, and sample concentrations were calculated based on this calibration curve. *p*-values were determined by one-way ANOVA followed by Tukey’s multiple comparisons test. The fitting of the Bradford assay standard curve, statistical analyses and data visualization were performed using Matlab R2023b (The Mathworks Inc.).

### qRT-PCR analysis

For real time quantitative PCR (qRT-PCR) analysis of *floA* mRNA in *A. fumigatus*, the wild-type strain A1160^+^ and mutant strain Δ*AFUB_007280* (*crzA*) were cultured in AMM for 20 h and then transferred to 0.2 M CaCl_2_ for 3 h. Mycelium was ground with a precooled mortar and pestle and total RNA was isolated with the Universal RNA Kit (Roboklon). Remaining DNA was digested with the TURBO DNA-free Kit (Thermo Fisher Scientific). All qRT-PCR reactions were performed on a CFX Duet Real-Time PCR System (BioRad), using the Luna Universal qPCR Master Mix (New England Biolabs). Three biological replicates for every strain were measured in three technical replicates each. Target gene expression was normalized to the expression of the reference gene β-actin using the 2 *^−^*^ΔΔ*Ct*^ method [91] and compared with A1160^+^ wild-type in AMM as a calibrator. Primer pairs for *floA* (*AFUB_024180*) and *β-actin* (AFUB_*093550*) were Afu_floA.FOR/Afu_floA.REV and Afu_actA.FOR/Afu_actA.REV, respectively. Both primer pairs had efficiencies between 90% and 110%.

### Detergent resistant membrane (DRM) isolation and Western blot analysis

For Western blot analyses, mycelia were homogenized in radioimmunoprecipitation assay (RIPA) buffer using a FastPrep homogenizer. Proteins were separated by SDS-PAGE using NuPAGE 4 to 12% (w/v) bis-Tris protein gels (Thermo Fisher Scientific) and transferred to a polyvinylidene difluoride (PVDF) membrane using iBlot3 PVDF ministacks (Thermo Fisher Scientific). The membrane was blocked with 5% (w/v) milk powder in TBS with 1% (v/v) Tween20 at room temperature for 1 h and then incubated overnight at 4 °C with the primary antibody. As primary antibodies, mouse HA Tag monoclonal antibody (Proteintech), GFP tag polyclonal antibody (Proteintech), mCherry polyclonal antibody (Proteintech) and GAPDH monoclonal antibody (Proteintech) as a loading control were used with a dilution of 1:1000 (GFP and mCherry), 1:2000 (GAPDH) or 1:50000 (HA). Hybridization of primary antibody with an HRP-linked anti-mouse IgG (Cell Signaling Technology) or HRP-linked anti-rabbit IgG (Abcam) was performed for 1 h at room temperature. Chemiluminescence of the HRP substrate (Merck Chemicals) was detected with a Fusion FX7 system (Vilber Lourmat).

For analysis of miniTurboID biotinylation activity, the strains *floA-3×HA-miniTurboID* and *3×HA-miniTurboID* were cultured in AMM for 18 h and then transferred to AMM with 0.1 M L-threonine and 0.1% (w/v) fructose as the sole carbon sources, supplemented with 50 µM biotin and cultured further for 3 h. After homogenization of mycelia in Pierce IP lysis buffer (Thermo Fisher Scientific), SDS-PAGE and Western blot were performed as described above, with the exception of using Western Blocking Reagent (Roche) instead of milk powder for blocking. The membrane was incubated with HRP-conjugated streptavidin (Thermo Fisher Scientific) at a 1:5000 dilution for 1 h at room temperature and then visualized as described above.

For the isolation of DRMs, *A. nidulans* strains *pmaA-mCherry*, *alcA*_p_*-floA-GFP* and SJW02 (GFP-TubA) [92] were cultured in AMM containing 0.1 M L-threonine and 0.1% (w/v) fructose in order to overexpress GFP-TubA and FloA-GFP under control of *alcA*_p_. The mycelium was harvested using Miracloth and resuspended in TNE lysis buffer (50 mM Tris-HCl pH 7.4, 150 mM NaCl, 5 mM EDTA) supplemented with a tablet of cOmplete Ultra Protease Inhibitor Cocktail (Roche) and centrifuged at 18,000 *× g* for 5 min at 4 °C. 125 µL of the supernatant was treated with 1% (v/v) Triton X-100 for 30 min on ice and then mixed with 250 µL Optiprep Density Gradient Medium (Sigma-Aldrich) to make a 40% (w/v) iodixanol solution. The sample was transferred to an ultracentrifuge tube and carefully overlaid with 600 µL of 30% (v/v) iodixanol in TNE lysis buffer and 100 µL TNE lysis buffer, then centrifuged at 100,000 *× g* for 2 h at 4 °C. Twelve fractions of equal volume were collected from the top of the gradient and analyzed by Western blot analysis.

### Proximity-dependent biotin identification (BioID) analysis

For BioID analysis, cell lysates of strains *floA-3×HA-miniTurboID* and *3×HA-miniTurboID* were prepared as described above for Western blot analysis of biotinylation activity with six biological replicates each. For enrichment of biotinylated proteins, 500 µL of cell lysate at 100 µg/mL protein concentration were rotated with 30 µL of Dynabeads MyOne Streptavidin T1 (Thermo Fisher Scientific) at 4 °C. The beads were then washed five times with PBS and the biotinylated proteins were eluted by resuspending the beads in 150 µL elution buffer (25 mM D-biotin, 2 % (w/v) SDS, 50 mM Tris HCl pH 8, 200 mM NaCl) and heating at 95 °C for 5 min. Cysteine thiols were reduced and carbamidomethylated in one step for 30 min at 70 °C by addition of each 4 µL of 500 mM TCEP (tris(2-carboxyethyl)phosphine) and 625 mM 2-chloroacetamide (CAA) per 150 µL eluates. Proteins were precipitated with MeOH/chloroform/H_2_O [93] and resolubilized in 100 mM TEAB in 5:95 trifluoroethanol/H_2_O (v/v). Proteins were digested for 18 h at 37 °C after addition of Trypsin/LysC protease mix (Promega). Tryptic peptides were dried in a vacuum concentrator and resolubilized in 200 µL 0.1% (v/v) trifluoroacetic acid (TFA). Peptides were cleaned up on C18 spin tips reconstituted with 200 µL ACN and equilibrated with 200 µL 0.1% (v/v) TFA. Enriched peptides were washed with 200 µL of 0.1% (v/v) TFA in 5:95 MeOH/H_2_O. Peptides were eluted by 200 µL 0.1% (v/v) TFA 80:20 ACN/H_2_O and evaporated to dryness. Finally, peptides were resolubilized in 30 µL of 0.05% (v/v) TFA in H_2_O/ACN 98/2 (v/v) filtered through 0.2 µm Ultrafree-MC hydrophilic PTFE filters (Merck Chemicals). The filtrate was transferred to HPLC vials and 4 µL were injected into the LC-MS/MS instrument.

Each sample was measured in three analytical replicates. LC-MS/MS analysis was performed on an Ultimate 3000 nano RSLC system connected to an Orbitrap Exploris 480 mass spectrometer (both Thermo Fisher Scientific, Waltham, MA, USA) with FAIMS. Peptide trapping for 5 min on an Acclaim Pep Map 100 column (2 cm *×* 75 µm, 3 µm) at 5 µL/min was followed by separation on an µPACneo 110 column. Mobile phase gradient elution of eluent A (0.1% (v/v) formic acid in water) mixed with eluent B (0.1% (v/v) formic acid in 90/10 acetonitrile/water) was performed using the following gradient: 0 min at 4% B and 750 nL/min, 10 min at 9% B and 750 nL/min, 12 min at 9.5% B and 300 nL/min, 55 min at 25% B and 300 nL/min, 70 min at 50% B and 300 nL/min, 75 min at 96% B and 300 nL/min, 78-80 min at 96% B and 750 nL/min, 80.1-90 min at 4% B and 750 nL/min. Positively charged ions were generated at spray voltage of 2.2 kV using a stainless steel emitter attached to the Nanospray Flex Ion Source (Thermo Fisher Scientific). The quadrupole/orbitrap instrument was operated in Full MS / data-dependent MS2 mode. Precursor ions were monitored at m/z 300–1100 at a resolution of 120,000 FWHM (full width at half maximum) using a maximum injection time (ITmax) of 50 ms and 300% normalized AGC (automatic gain control) target. Precursor ions with a charge state of z=2–5 were filtered at an isolation width of m/z 4.0 amu for further fragmentation at 28% HCD collision energy. MS2 ions were scanned at 15,000 FWHM (ITmax=40 ms, AGC=200%). Each sample was measured in triplicate with a different compensation voltage (-42 V, -57 V, -72 V).

Tandem mass spectra were searched against the *A. nidulans* UniProt proteome databases (2024/06/03 (YYYY/MM/DD) (https://www.uniprot.org/proteomes/UP000000560) using Proteome Discoverer (PD) 3.1 (Thermo) and the database search algorithms Mascot 2.8, Comet, MS Amanda 2.0, Sequest HT with and without INFERYS Rescoring, and CHIMERYS. Two missed cleavages were allowed for the tryptic digestion. The precursor mass tolerance was set to 10 ppm and the fragment mass tolerance was set to 0.02 Da. Modifications were defined as dynamic Lys biotinylation, Met oxidation, protein N-term acetylation with and without Met-loss as well as static Cys carbamidomethylation. A strict false discovery rate (FDR) *<*1% (peptide and protein level) was required for positive protein hits. Furthermore, search engine scores were filtered for either Mascot (*>*30), Comet (*>*3), MS Amanda 2.0 (*>*300), Sequest HT (*>*3) or CHIMERYS (*>*2). The Percolator node of PD3.1 and a reverse decoy database was used for *q*-value validation of spectral matches. Only rank 1 proteins and peptides of the top scored proteins were counted. Label-free protein quantification was based on the Minora algorithm of PD3.0 using the precursor abundance based on intensity and a signal-to-noise ratio *>*5. Normalization was performed by using the total peptide amount method.

Imputation of missing quan values was applied by using abundance values of 75% of the lowest abundance identified. Differential protein abundance was defined as a fold change of *>*2, *p*-value/ABS(log4ratio) *<*0.05 and at least identified in five of six biological replicates of the sample group with the highest abundance. Statistics was performed with R 4.4.0 and RStudio 2024.04.0. Data visualization was performed with Matlab R2023b (The Mathworks Inc.).

### Light microscopy

For visualization with differential interference contrast (DIC) microscopy, conidia were incubated on coverslips in AMM at 25 °C for 20 h. For fluorescence microscopy, conidia were incubated overnight in glass-bottom ibi-Treat 8-well chamber slides (ibidi) in AMM for 20 h at 25 °C. For FloA-GFP expression regulated by *alcA*_p_, conidia were grown in AMM with 0.1% (w/v) fructose and 0.1 M L-threonine as the carbon source. Calcium stress was applied by exchanging the medium to 0.2 M CaCl_2_ and further incubation for 3 h at 37 °C. Ionic stress and osmotic stress were applied by exchanging the medium to 0.2 M KCl, 0.2 M NaCl, 0.2 M MgCl_2_ or 0.6 M sorbitol. Cell viability was evaluated by staining with 1 µM Sytox Blue (Thermo Fisher Scientific) for 10 min. Intracellular calcium was visualized using the Fluo-4 Direct Calcium Assay Kit (Thermo Fisher Scientific) according to the manufacturer’s instructions. Samples were visualized using a Zeiss LSM 780 confocal microscope and images were processed with Fiji [94]. The intensity profile along the cytoplasmic membrane of the conidium was analyzed using the oval profile plugin [95] in Fiji, after cutting out all signals inside the cell. For each channel, the intensities were normalized by the maximum intensity and plotted using Matlab R2023b (The Mathworks Inc.).

CellTracker Blue CMAC (Thermo Fisher Scientific) was used at a final concentration of 10 µM in medium from a stock solution of 10 mM in DMSO, and incubated at 37 °C for 15 min. The staining was followed by two washing steps and resuspension in 0.9% (w/v) NaCl. Fluorescence and bright-field images were taken using a Keyence BZ-X800 microscope (Keyence). For each fluorescence image, the background was subtracted using a rolling ball algorithm (radius: 50 pixels) on Fiji [94].

## Acknowledgements

We are grateful to Christina Täumer for excellent technical assistance and Lei-jie Jia for providing plasmid pLJ-HscA-mT. Stefanie Pöggeler and Lucas Hollstein are appreciated for their advice on BioID analyses in filamentous fungi. M.K. received funding from the Deutsche Forschungsgemeinschaft (DFG) (www.dfg.de) Cluster of Excellence “Balance of the Microverse” (EXC 2051; project-ID 390713860). T.K. and M.R. were funded by DFG CRC/Transregio 124 “FungiNet” (project number 210879364; projects A1, Z2) and DFG CRC 1127 “ChemBioSys” (project number 239748522; project B02), respectively. M.K. and S.T.G. were also supported by the DFG Excellence Graduate School “Jena School for Microbial Communication”.

## Supporting information

Supplementary data are included in a separate file.

## Supporting information captions

**S1 Table.** Strains used in this study

**S2 Table.** List of primers used in this study

**S3 Appendix.** Generation of *A. nidulans* mutant strains

**S4 Fig.**: Deletion of the *floA* gene in *A. nidulans*. (a) Strategy for *floA* deletion by double cross-over leading to the replacement of *floA* against *argB* on the chromosome. The blue, green, red and yellow arrows depict the binding sites of the primer pairs used for the PCR verification in (b), (c), (d) and (e), respectively. (b) PCR using primers floA.FOR/floA.REV. Presence of *floA* is indicated by a 1.6 kb band. (c) PCR using primers floA up.FOR/floA down.REV. Presence and absence of *floA* are indicated by 2 kb and 1.8 kb bands, respectively. (d) PCR using primers floA 5HOM1kb^+^.FOR/argB CDS.REV. Presence of the *argB* cassette is indicated by a 2.4 kb band. (d) PCR using primers argB CDS.FOR/floA 3HOM1kb^+^.REV. Presence of the *argB* cassette is indicated by a 2.6 kb band.

**S5 Fig.**: Localization of PmaA-mCherry in the _Δ_*floA* background in 0.2 M CaCl_2_, 0.2 M KCl, 0.2 M NaCl and 0.6 M sorbitol.

## Accession numbers

The mass spectrometry proteomics data have been deposited to the ProteomeXchange Consortium via the PRIDE [96] partner repository with the dataset identifier PXD076500 and 10.6019/PXD076500.

